# Multi-omics Analysis Reveals Important Role for Microbial-derived Metabolites from *Botryllus schlosseri* in Metal Interactions

**DOI:** 10.1101/2024.11.10.622856

**Authors:** Dulce G. Guillén Matus, Caroline M. Donaghy, Nidhi Vijayan, Zachary T. Lane, Matthew Howell, George G. Glavin, Alfredo M. Angeles-Boza, Spencer V. Nyholm, Marcy J. Balunas

## Abstract

Marine microbial communities govern many of the biological and chemical processes in the ocean, including element cycles, ecosystem health, and disease. Marine organisms are surrounded by microbes and complex molecular interactions occur between bacterial symbionts, eukaryotic hosts, and their pathogens or prey. Trace metals in the ocean can be either beneficial or detrimental to marine life depending on their concentrations and bioavailability. Multiple marine tunicate species are known to bioaccumulate trace metals in their mantel, and research suggests tunicate microbiota plays an important role in this process. *Botryllus schlosseri*, a marine colonial tunicate, has become a model organism for cellular and developmental studies, yet its ecological interactions are still not well understood. Using an integrated multidisciplinary approach, we established a comprehensive baseline and explored correlations between members of the *B. schlosseri* microbiome, metabolome, and metallome to elucidate the ecological effects of trace metals in host-microbe-pathogen interactions. We identified significant correlations between metals, including manganese, nickel, cerium, zinc, and cobalt, with various metabolites and bacterial taxa. These findings offer insights into *B. schlosseri* biological and chemical interactions with their symbionts and their environment, contributing to bridging the knowledge gap of host-microbiome-environment interactions and establishing a foundation for continuing research on the ecological effects of trace metals in these biological systems.

**Graphical Abstract:** 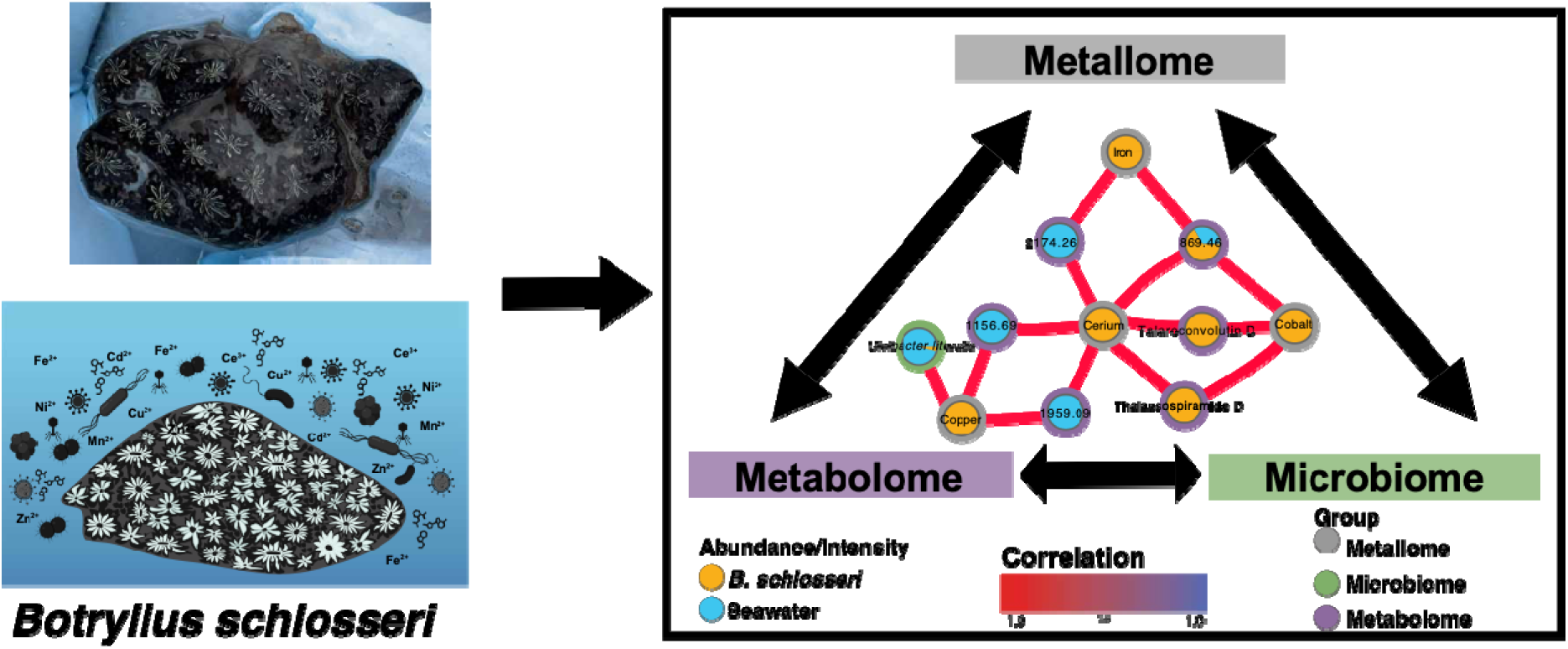

**Highlights:**

- *Botryllus schlosseri* tissue was highly enriched in metals compared to seawater
- *B. schlosseri* microbiome ß-diversity significantly different from seawater
- Pan-metabolome indicated microbial metabolites in core and flexible metabolome
- Multi-omics revealed interactions between metals, metabolites, and microbes

## 1. Introduction

Metal ions in marine environments play a pivotal role, having both positive and negative effects to marine ecosystems while influencing ecological processes and contributing to the overall health of these ecosystems. About a dozen elements with atomic mass above 50 are known to have biological roles, often as cofactors, promoters of redox chemistry, or as structural elements in proteins (1, 2). Iron, manganese, nickel, zinc, cobalt, and vanadium have been the most characterized in oceanography (3). Iron, probably the most well studied, is a limiting nutrient for most biological processes and plays an essential role in primary production, typically found in planktonic communities at high concentrations (3, 4). However, although the biochemical functions of many metal ions are extensive, we currently lack a complete understanding of the role played by these important trace elements, especially in complex marine environments (3, 5). It has been demonstrated that small changes in metal concentrations, as low as sub-nanomolar, can have drastic effects in marine environments (6). Coastal environments, enriched in many trace metals, exhibit a remarkable biological diversity, in part facilitated by the interface between terrestrial and oceanic influences, fostering a unique and vital habitat for a wide array of marine species of microbes and complex organisms (7, 8).

Microbial communities in the ocean govern many processes including primary production, ecosystem health and disease, and many element cycles (9). They are found in planktonic states in the water column, forming biofilms on solid surfaces, and/or inhabiting macroorganisms (10). Interactions between microorganisms and animals can range from the broad categories of mutualism, in which both parties’ benefit, to pathogenic, in which interactions harm one or more parties. Marine invertebrates, especially filter feeders, comprise the largest biodiversity of multicellular eukaryotes in the ocean, and harbor complex relationships with microbial communities (11, 12). Many marine invertebrates enter into beneficial microbial symbioses that allow these species to take advantage of habitats that would otherwise be unavailable to them (e.g., tropical corals and hydrothermal vent tubeworms; (13–15).

The study of host-associated microbial communities is crucial for the understanding of animal physiology, especially as related to pivotal roles for microbiota in host development, nutrition, and behavior (16, 17). However, more recently microbiome research has often been paired with metabolomics analyses to further understand how host chemical output shapes their microbiota and how microbially-derived molecules influence host physiology (17, 18). While genomics (e.g., 16S rRNA community profiles) provide insight into the microbial taxa present in the host, metabolomics provides a chemical fingerprint of cellular processes occurring within and between hosts and microbes (19, 20). Current comparative metabolomics methods allow for rapid analysis of large number of samples, facilitating chemical ecology investigations across host species and ecosystems, including their complex associations with microbial communities.

Tunicates, classified under the phylum Chordata, are filter-feeding invertebrates known to harbor bacteria in their digestive tracts, tunics (21), and throughout their bodies (22). In some cases, tunicate-associated bacteria have been shown to be symbiotic or mutualistic with the host, such as defending against pathogenic microbes, acting as feeding deterrents, and/or helping with nutrient acquisition, thus making them vital for host survival (23, 24). Tunicates have also emerged as noteworthy sources of natural products, known for their valuable metabolites with antimicrobial, antiparasitic, and anticancer properties (25, 26, 24, 27). Tunicates are also important for marine food webs, triggering biogeochemical flux from the surface to deep waters, and have even been linked to being potential environmental stress indicators (28), including as bioindicators for metals in marine ecosystems (29, 1, 30).

The colonial tunicate, *Botryllus schlosseri*, is a cosmopolitan species that inhabits shallow intertidal marine habitats around the world (31). *B. schlosseri* has emerged as a prominent model organism in many areas of biology, including questions of histocompatibility, self/non-self recognition, stem cell biology, and development (32–35). However, little is known about the microbiome of *B. schlosseri* with even more limited studies of the metallome or metabolome of this species. Two microbiome studies of this and other tunicates found their microbial communities to be conserved within species, suggesting that the mucosal layer of the pharynx may harbor specific symbionts (31, 36). Thus, although there has been emphasis on developing *B. schlosseri* as a model organism, there are still significant gaps in the understanding of its ecology and environmental adaptations.

Using a multidisciplinary approach, we investigated the *B. schlosseri* microbiome, metabolome, and metallome, including multi-omics integration to determine the importance of microbes, metabolites, and metals in *B. schlosseri* communities. Our integrative multi-omics approach provides a novel perspective regarding how these tunicates flourish in their environment, aimed at further elucidation of the importance of microbes, metabolites, and metals in these marine systems. Our findings provide the first steps for understanding ecological effects of trace metals in the host-microbe interactions of *B. schlosseri.* Studying the intricate relationship between *B. schlosseri*’s microbiome, metabolome, and metallome helps to unravel the complex interplay between these organisms and their environment, providing crucial insights into the ecological dynamics, adaptive strategies, and potential bioaccumulation effects in marine ecosystems.

## 2. Experimental section

### 2.1 Tunicate and seawater sample collection

Eight subtidal *B. schlosseri* colonies, identified via phenotypical characteristics, were collected from artificial substrates via submerged ropes and other structures from three dock sites at Avery Point in Groton, CT (N 41°18’59”, W 72°3’39”). *B. schlosseri* colonies were quickly dipped in 100% ethanol (EtOH) to surface sterilize, rinsed with sterile artificial seawater (40g/L Instant Ocean^®^ Sea Salts, Blacksburg, VA) 2-3 times and separated into three aliquots, one for each type of analysis. Samples were then placed on ice and transported to the lab where they were stored at -80 °C until further processing.

Surface seawater was collected from the same three tunicate collection sites in 5L quantities. Seawater collections were brought back to lab and processed immediately. For microbiome and metabolomic analyses, 1L from each collection was vacuum filtered through a sterile vacuum apparatus (Millipore^®^) equipped with 0.22µm, 47 mm diameter, autoclaved PTFE hydrophobic filter disks (Millipore^®^), repeated in triplicate to obtain samples for each type of analysis. Filter disks were removed from the apparatus and stored at -80 °C until analysis. For metals analyses, 1L volume of seawater was transferred to ICP digiprep tubes (SCP Science, DigiTUBES) and kept in a cooler during transportation to the lab. The seawater was filtered with a sterile 0.22 μm filter (ThermoScientific, Nalgene Prefilter Plus), preserved with trace metal grade HCl, and stored at -10 °C until analysis.

### 2.2 Metallomics

#### 2.2.1 Metal quantification using ICP

Metal analyses were conducted using *B. schlosseri* or seawater samples. Quality control samples (i.e., laboratory reagent blank, laboratory fortified blank, laboratory fortified matrix, standard reference matrix, and calibration standards) were utilized for all experiments, prepared using the same matrices of digestion as each sample. Calibration standards for tunicate samples were prepared using the multi-element standard solution ICP Stock Standard and ICP-MS Memory Check Solution B (High Purity Standards, Charleston, SC). For seawater samples, calibration standards were prepared using the multi-element standard solution QC1 (Inorganic Ventures Labs, Christiansburg, VA). The acid matrix for digestion of tunicate samples used NanoPure water (filtered using a Barnstead NANOpure Diamond filtration system with a 0.2 μm pore size filter), HNO_3_, and H_2_O_2_. The acid matrix for digestion of the seawater samples used HNO_3_ and HCl. The quality controls and calibration standards for seawater samples used Instant Ocean (Instant Ocean, Blacksburg, VA) which was prepared to a concentration of 16.9089 g/L with NanoPure water. Analytical precision was measured as relative standard deviation which was consistently below 5%.

#### 2.2.2 Metal analysis of B. schlosseri samples

Prior to digestion, samples were lyophilized using a FreeZone Freeze Dryer (Labconco Corporation, Kansas City, MO) overnight and homogenized using an agate mortar and pestle. Most samples were analyzed individually with the exception of the “mixed” sample which consisted of four individual tunicate samples that were homogenized together in order to evaluate metal concentration yields from a larger sample. Tunicates were digested following modified EPA protocols 200.3 (37). Each sample was refluxed in trace metal grade HNO_3_ for 1 hour using a hot block (Environmental Express, Charleston, SC) at 92-95 °C. Once samples were cooled, 0.3 mL of NanoPure water and 0.75 mL of trace metal grade H_2_O_2_ were added, and samples were reheated until bubbling subsided. Samples were then cooled and brought to a final volume of 15 mL with NanoPure water. Filtering was unnecessary due to complete digestion, as indicated by lack of precipitants in solution. Samples were analyzed using inductively coupled plasma mass spectrometry (ICP-MS; Perkin-Elmer ELAN/DRC-e, Shelton, CT) following EPA analysis method 6020A (38) to determine metal concentrations of vanadium (V), manganese (Mn), iron (Fe), cobalt (Co), nickel (Ni), copper (Cu), zinc (Zn), and cerium (Ce). The limits of detection (LODs) were determined to be below 0.2 ppm for each metal, except for iron and zinc which had LODs below 15 and 3 ppm, respectively (Table S1).

#### 2.2.3 Metal analysis of seawater samples

Seawater samples were filtered with a 0.20 μm Nalgene Prefilter Plus GFP+ 0.2 μm CA filter (Thermo Scientific, Waltham, MA), preserved with trace metal grade HNO_3_, and stored at 4 °C until digestion. Seawater samples were digested following EPA protocols 200.7 (39). Preserved seawater samples were individually transferred to new digestion tubes in 25 mL increments to which 0.25 mL trace metal grade HNO_3_ and 0.125 mL of trace metal grade HCl was added and refluxed at 92-95 °C within a hot block for 2 hours or until samples had reduced in volume by 12 mL. Samples were then cooled and brought to the final volume of 25 mL using NanoPure water. Processed seawater samples were analyzed using an ICP-Optical Emission Spectrometer (ICP-OES; Spectro Arcos ICP-OES, SPECTRO Analytical Instruments GmbH, Kleve, Germany) to determine metal concentrations of V, Mn, Fe, Co, Ni, Cu, Zn, and Ce via true axial plasma observation. The LODs for metals in the seawater samples were determined to be equal to or less than 0.002 ppm (Table S1).

#### 2.2.4 Statistical analysis

Bioaccumulation results for tunicate samples were calculated via fold increase analysis, dividing the average metal value within each tunicate sample by the average metal value found in the surrounding seawater. Error corresponding to metal accumulation was calculated using error propagation since the *B. schlosseri* and seawater samples were derived from different populations. After averaging individual metal concentrations for both *B. schlosseri* and seawater samples, the corresponding average and standard deviation values were applied to Equation 1, where *z* corresponds to accumulation value, while *x* and *y* represent the average water and tunicate samples. All statistical analysis for the metal data was processed in Microsoft Excel using two-tailed, homoscedastic t-test analysis.

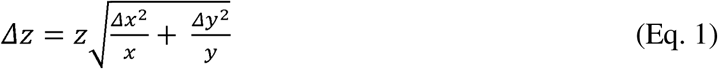

### 2.3 Bacterial community analysis

#### 2.3.1 DNA extraction and 16S rRNA analysis

Tunicate samples and seawater filters were thawed, bead-beated with Fastprep-24^TM^ (MP Biomedicals, Santa Ana, CA) at 4.5m/s for 45 s, centrifuged at 16,000 x g for 1min, and the supernatants was subjected to DNA extractions. Two replicates of negative controls, i.e. tubes without tissues, were processed alongside all samples. DNA was extracted using Qiagen PowerMag soil DNA isolation kits (27100-4-EP). The V4 region of the 16S rRNA gene was amplified via PCR using primers 515F (GTGYCAGCMGCCGCGGTAA) and 806R (GGACTACNVGGGTWTCTAAT). PCR conditions included an initial denaturation at 95 °C for 3 min, followed by 30 cycles of 95 °C for 30 s, 55 °C for 30 s, and 72 °C for 1 min and 30 s, with a final extension at 72°C for 10 min. PCR products were quantified using Qubit, purified with the QIAquick PCR Purification Kit, and sequenced using the Illumina MiSeq platform with the MiSeq Reagent Kit v2 (2 × 250 bp) at the University of Connecticut Microbial Analysis, Resources, and Services (MARS) facility. Sequences were deposited in the National Center for Biotechnology Information (NCBI) Sequence Read Archive (SRA) (accession number: PRJNA1039848).

Raw sequences were denoised and demultiplexed using QIIME 2.2020 (40) and amplicon sequence variants (ASVs) were generated using DADA2 (41). Sequences were aligned against a Naïve-Bayes trained Greengenes reference database (V2.2022) at a 99% or higher identity to the reference sequences (42). Sequences assigned as chloroplast or unassigned at Kingdom level were removed. Data processed with QIIME2 were converted to phyloseq objects using *qiime2R* (v0.99.6). The final ASV-abundance matrix was normalized with *transform_sample_counts* to the minimum number of reads in the sample set (6933). We also rarefied the data using *rarefy_even_depth* and compared the statistical data with normalization and found both methods generated similar results. Beta diversities were obtained using the Bray-Curtis (43) and Weighted UniFrac (44) modules in QIIME2 and plotted in R using the phyloseq package. Alpha diversities, including Observed, Shannon, and Inverse Simpson indices, were calculated with *estimate_richness* function from the phyloseq package.

### 2.4 Metabolomics

#### 2.4.1 General experimental

All solvents for LC-MS/MS were LC-MS grade from Sigma-Aldrich (St. Louis, MO). Solvents for extraction and semi-preparative high-performance liquid chromatography (HPLC) purification were HPLC grade from Sigma-Aldrich (St. Louis, MO).

#### 2.4.2 B. schlosseri and seawater extraction for mass spectrometry (MS) metabolomics

Tunicate samples or seawater filters were thawed at room temperature and all extraction glassware, containers, and tools were pre-rinsed with extraction solvents. Samples were transferred to glass beakers and extracted four times with 2:1 dicloromethane:methanol (DCM:MeOH; ∼5-10mL), including maceration, stirring, and filtering to afford an exhaustive extraction. Extracts were then concentrated via rotary evaporation at 38 °C using a Büchi rotavapor model R-210 (New Castle, DE, USA), transferred to vials, and stored at -80 °C.

#### 2.4.3 Mass spectrometry (MS) data acquisition and processing

Extracts were prepared at 1 mg/mL in methanol (MeOH). Data was acquired using a Bruker timsTOF Pro2 coupled to an ELUTE UPLC (Bruker-Daltonics, Billerica, MA, USA). Each sample (2 μL) was injected in technical triplicate using a Bruker Intensity Solo 2 C_18_ column (100 x 2.1 mm, 1.8 μm). Samples were eluted using a 0.3 mL/min gradient of mobile phases A (0.1% formic acid in water [H_2_O]) and B (0.1% formic acid in acetonitrile [ACN]) beginning at 5% B for 0.5 min, followed by a linear gradient from 5 to 40% B for 3.5 min, then 40 to 98% B for 4.0 min, holding at 98% B for 1.0 min, followed by a return to 5% B for 0.5 min, holding at 0.5% B for 1.0 minute.

Data acquisition was performed in positive ionization mode using a VIP-HESI (Vacuum Insulated Probe Heated Electrospray Ionization) source, with a collision energy of 10 eV, capillary voltage of 4500 V, dry temperature of 220 °C, sheath gas temperature of 400 °C, mass range of 150-2200 *m/z*, and mobility (1/K_0_) range of 0.55–1.90 V·s/cm^2^, and ramp time of 100 ms. Fragmentation data were acquired with a collision energy of 50 eV, with 2 PASEF MS/MS scans per cycle, active exclusion release after 0.1 min, for a total cycle of 0.53 seconds.

Once acquired, MS data were preprocessed using Bruker MetaboScape^®^ version 9.0.1 (Bruker-Daltonics, Billerica, MA, USA) using the MCube T-Rex 4D Metabolomics workflow for peak picking and alignment. The intensity threshold was experimentally determined by comparison of baseline noise from samples and MeOH blanks resulting in an intensity threshold of 7500 counts, resulting in a feature table with 2545 total features before filtering.

The resulting feature table was then processed using the MPACT software (45) with the following parameters: mispicked peak correction – ringing mass window of 0.5 atomic mass units (AMUs), isotope mass window of 0.01 AMU with a maximum isotopic mass shift of 3 AMUs, and a *t_R_* window of 0.05; in-source ion filtering threshold of 0.95 Spearman correlation; median coefficient of variation (CV) of technical replicates of 0.5; and blank filtering using MeOH blanks using a 0.05 threshold. After MPACT filtering, 17.3 % of features passed the filtering steps, removing 1243 blank features, 217 mispicked features, and 281 features found to be nonreproducible across technical replicates.

For *in silico* formula prediction and annotations, MetaboScape^®^, MPACT (45), Sirius (46), NPAtlas (47), and MarinLit (48) were used as databases for annotations, with a 10 ppm upper limit for both annotation and formula prediction. When pertinent, annotations were further verified by comparing fragmentation patterns using public data, when available, or using the Competitive Fragmentation Modeling for Metabolite Identification (CFM-ID) spectra prediction (49). Confidence for each annotation was assessed using Schymanski’s rules for metabolite annotation [Table S5; (50)].

#### 2.4.4 Metabolomics and multivariate analysis

MPACT (45) was used to conduct statistical comparisons between metabolomes of *B. schlosseri* and seawater, using heatmaps, non-metric multidimensional scaling (NMDS) analyses, and volcano plots. The heatmap was built with normalized raw counts within each feature between minimum and maximum counts for that feature. Samples were grouped on the x-axis by overall metabolomic similarity, with features grouped on the y-axis. The NMDS plot was generated using Bray-Curtis dissimilarity distance, after averaging technical replicates. The volcano plot was generated using multiple two tailed t-tests, including false discovery rate (FDR) correction using the Benjamini-Hochberg procedure (51).

For construction of the *B. schlosseri* pan-metabolome, the presence or absence of all features was calculated for every individual. Using group parsing calculations (e.g., (52, 45) a feature was present if the ratio between its maximum count across all *B. sclosseri* samples to the count from each individual was below 100. A feature was considered absent if its counts were zero or if the maximum over the individual ratio was 100 or higher. To visualize the *B. schlosseri* pan-metabolome, an upset plot was generated using the R package UpsetR (53), adding stacked bars to indicate unannotated features as well as the putative origin of any annotated metabolites according to the information provided in databases used for annotation, classifying each feature as from microbial, invertebrate, or mixed origin, when reported from both microbes and invertebrates.

### 2.5 Multi-omics integration model

Integration of the metabolome, microbiome, and metallome was accomplished using Diablo [Data Integration Analysis for Biomarker discovery using Latent variable approaches for Omics studies; (54)], which is an N-integration model from the mixOmics R package (55). A subset of three *B. schlosseri* colonies for which data across all -omics were available were utilized for multi-omics integration analysis. Comparison of metals concentrations, OTUs sequence counts, and MS data, was facilitated by scaling each data type, beginning with scaling each metal concentration across its own maximum and minimum. Metabolites were normalized using square root and used pareto scale (56). The OTU sequence counts were normalized using center log ratio (57). Further, any feature with a standard deviation of zero across all data was removed, since these would not contribute to the Diablo covariance model.

To integrate multi-omics data, the N-integration model in Diablo can be run using either full or tuned mode to generate the model. The tuned mode (reduced model) was used for generation of a circos plot based on Spearman’s pairwise correlation coefficient (-1 ≤ r ≥ 1) and included only the first and second component of variance of the integration model to reduce complexity for easier visualization. To choose the optimal number of components, an M-fold validation was performed with 3 folds and 10 iterations, with the number of components to be tested set at 3. Similarly, the optimal number of features per -omic dataset for each of the components (from the previous step) was determined by running an M-fold validation with 3 folds and 10 iterations. Lastly, a hyperparameter of 0.1 was used for the design matrix. The full integration model was used to create a multiblock sparse partial least squares discriminant analysis [(s)PLS-DA] (Figs. S5 and S6) and a correlation network with a hyperparameter of 0.1 for the design matrix and three components. Subsequently, network percolation was performed using the R package *associationSubgraph* (58) to set up correlation thresholds to facilitate prioritization of network interactions and allow for easier visualization of correlation clusters. The resulting thresholds for the connections were 0.995 for positive connections and 0.990 for negative connections. Networks were visualized using Cytoscape (59).

## 3. Results and discussion

### 3.1 Metallomics of B. schlosseri and surrounding seawater

Using acid digestion and ICP analysis, the average concentrations of vanadium (V), manganese (Mn), iron (Fe), cobalt (Co), nickel (Ni), copper (Cu), zinc (Zn), and cerium (Ce) were determined for both *B. schlosseri* tissues as well as their surrounding seawater (Fig. 1, Table S2). The concentrations of all measured metals varied relative to one another (Fig. 1A). Iron, manganese, and zinc were the most abundant metals in *B. schlosseri*, while cobalt was the least abundant metal found in the tunicates. The rare earth metal, cerium was also found to be highly abundant in *B. schlosseri*. In seawater, iron, nickel, and manganese were the most abundant, while cobalt and cerium were the least abundant. To determine the extent to which these metals were accumulated in *B. schlosseri* relative to their environment, we also established fold increases for each metal (Fig. 1B). Concentrations of all measured metals were found to be significantly (p < 0.01) higher in *B. schlosseri* than in seawater, especially zinc, manganese, and iron, indicative of tunicate metal sequestration.

**Fig. 1.**
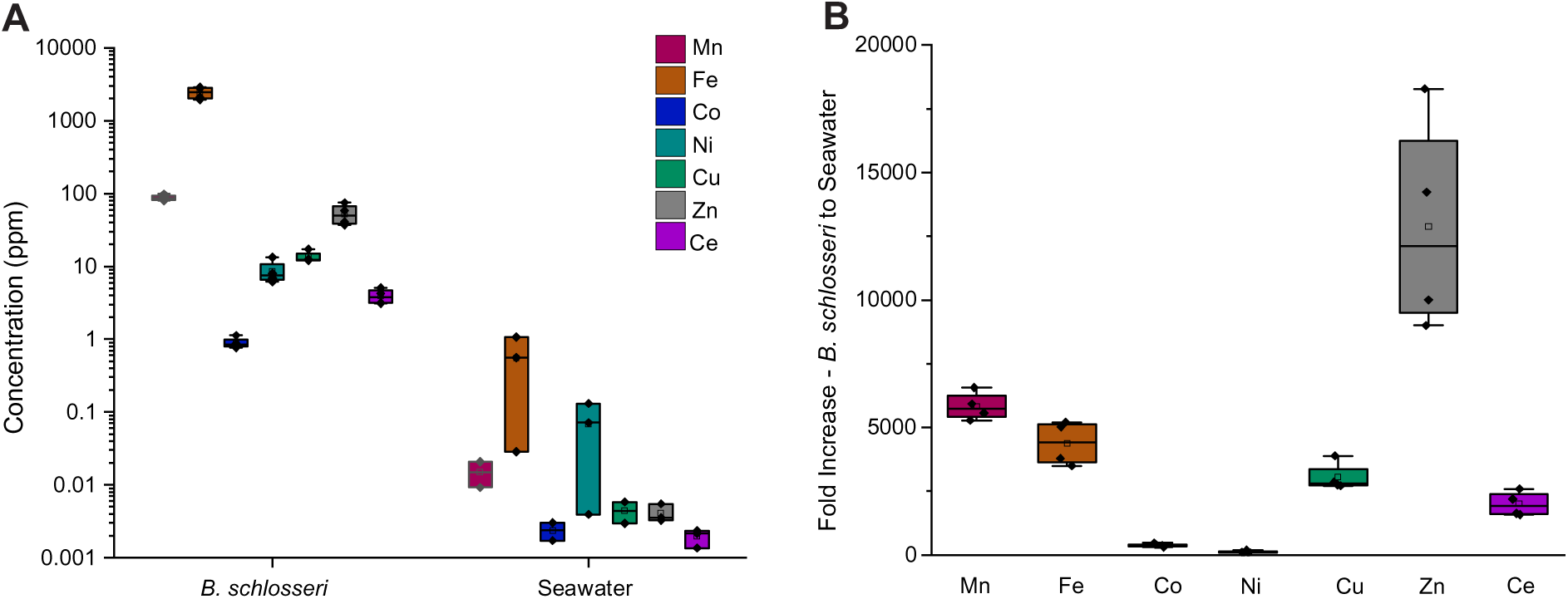
*B. schlosseri* tunicates sequester metals at substantially higher concentrations compared with surrounding seawater. (A) Average metal concentrations for *B. schlosseri* and seawater samples in units of parts per million [y-axis on logarithmic scale; vanadium (V), manganese (Mn), iron (Fe), cobalt (Co), nickel (Ni), copper (Cu), zinc (Zn), and cerium (Ce)], indicating substantially higher concentrations of all metals within *B. schlosseri*. Vanadium data not shown for seawater because concentrations were below limit of detection (LOD). (B) Fold increase of average *B. schlosseri* metal concentrations relative to those of the surrounding seawater (error bars represent the variation in the fold increases across samples including error propagation).

Our findings are consistent with previous well-established research on the natural abundance of many of these metals, e.g. iron, manganese, zinc, and copper, in seawater (61–64) and their role as common cofactors in marine biochemistry (65, 66). Zinc was the most substantially increased in *B. schlosseri* versus seawater, which is not surprising given its important role in the structure and function of enzymes involved in gene expression, protein synthesis, and immune function (67, 68). Manganese was the second-most bioaccumulated metal in our study and is known for its role in energy metabolism and hormone synthesis (69, 70). Iron was also heavily sequestered by *B. schlosseri* and is crucial for many biochemical reactions in biological systems, but is also used for electron transport, DNA synthesis, and regulation of both gene expression and cellular signaling (71, 72). Interestingly, cerium was also bioaccumulated, especially when considering how little cerium was present in the surrounding seawater. Previous studies have found tunicates sequester substantial quantities of rare earth metals, including cerium (45). This accumulation may be indicative of the tunicates may exploit unique biochemistry for their proliferation and invasion of habitats. Although the role of cerium has not yet been elucidated for *B. schlosseri*, cerium may aid in antioxidant activity, redox chemistry, detoxification, or structural enhancement of the tunicate tissue (73, 74). Copper was also sequestered by *B. schlosseri*, with previous work finding that copper accumulation may aid in deterrence of predators due the delicate balance between accumulation and toxicity for both tunicates and their larvae (75). The least bioaccumulated metals in *B. schlosseri*, nickel and cobalt, are both heavy metals that have low thresholds for toxicity but that are often found to be structural components within enzymes and proteins of biological systems. Although the roles for these metals are not yet defined in *B. schlosseri,* our research provides possible insights into their importance for this colonial tunicate.

### 3.2 Bacterial diversity of B. schlosseri and surrounding seawater

To generate 16S rRNA community profiles, microbial DNA from eight *B. schlosseri* and three seawater samples was amplified and sequenced, resulting in total amplicon reads of 276,709 and 196,968, respectively. The number of amplicon reads for *B. schlosseri* and seawater samples ranged from 4173-81,886 and 61,546-69,946, respectively. After normalizing reads from *B. schlosseri* samples to the minimum number of reads, *B schlosseri* communities were shown to have 2200 unique taxa, whereas the normalized dataset for seawater samples had 2571 taxa. Plateauing of the rarefaction curves was observed for both *B. schlosseri* and seawater samples (Fig. S1).

To consider microbial diversity at the community level, alpha and beta diversity were calculated (Fig. 2A and 2B). Using the Wilcoxon test, no significant differences (p > 0.05) in alpha diversity were observed between *B. schlosseri* and seawater samples (Fig. 2A), possibly due to tunicates being filter feeders and animals were not depurated before sampling. However, significant differences in beta diversity between tunicate and seawater microbial communities were identified using Weighted Unifrac (PERMANOVA q-value = 0.001, pseudo-F = 11.82) and Bray-Curtis distance metrics (PERMANOVA q-value = 0.001, pseudo-F = 4.34). This larger effect size with Weighted Unifrac analysis suggests a stronger difference between microbiomes when we consider phylogenetic relatedness and the smaller effect size with Bray-Curtis suggests differences are less pronounced when we consider abundance alone, without considering evolutionary relationships (43, 44). Further, the differences observed can also be partially attributed to the within-sample variation between replicates of *B. schlosseri* (Weighted Unifrac PERMDISP p-value = 0.048), though this result is only marginally significant. This suggests that while tunicate-associated microbiomes may exhibit greater heterogeneity compared to seawater communities, additional factors likely contribute to the observed differences in beta diversity.

**Fig. 2.**
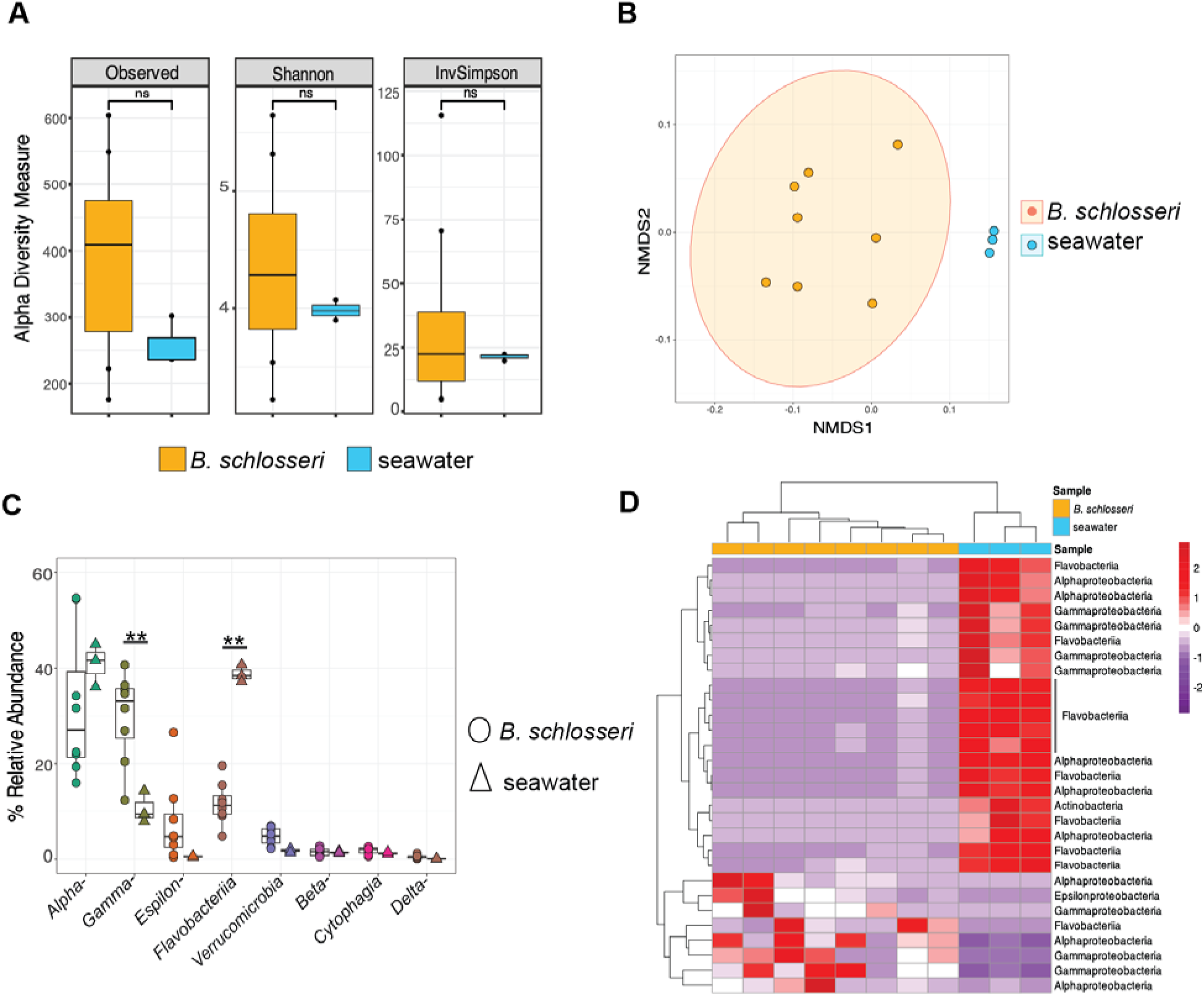
Bacterial diversity in *Botryllus schlosseri* and surrounding seawater. (A) No significance difference was found in alpha diversity between *B. schlosseri* and surrounding seawater (p > 0.05). (B) Non-metric multidimensional scaling (NMDS) of beta diversity metric, using weighted UniFrac, demonstrated the bacterial community composition of *B. schlosseri* was distinct from the surrounding seawater. (C) Percent relative abundance of bacterial taxa making up > 0.1% of the *B. schlosseri* and seawater microbiota indicated that some taxa are more abundant in the tunicate (t-test; p-value ** < 0.01). (D) Heatmap of the most differentially abundant amplicon sequence variants (ASVs) classified to their Class taxonomic level.

To further explore the microbial communities of *B. schlosseri* and surrounding seawater, relative abundances of specific taxa were compared (Fig. 2C and 2D). In *B. schlosseri*, the most abundant classes were Alphaproteobacteria (31 ± 12%), Gammaproteobacteria (29 ± 6.7%), and Flavobacteriia (11.7 ± 4%), comprising approximately 73% of the microbiota (Fig 2C). Other abundant taxa in *B. schlosseri* included Epsilonproteobacteria (7 ± 6%), Verrucomicrobia (4.7 ± 1.1%), Betaproteobacteria (1.54 ± 0.8%), Cytophagia (1.8 ± 0.4%) and Deltaproteobacteria (0.54 ± 0.34%), with diatoms comprising 7.5 ± 7.5% relative abundance (Fig. 2C, Fig. S2). Further analyses were conducted to determine microbial taxa that were enriched in *B. schlosseri* compared with surrounding seawater (Fig. 2D), finding enrichment in *B. schlosseri* of ASVs of Alphaproteobacteria, Gammaproteobacteria, Flavobacteriia, and Epsilonproteobacteria (Fig 2D). Many taxa identified in our *B. schlosseri* bacterial communities are commonly found in seawater and often associated with other marine organisms. For example, Alphaproteobacteria, Gammaproteobacteria, and Flavobacteriia were previously identified in the microbiota of *B. schlosseri* and other tunicates including *Cinoa robusta*, *C. savigni*, and *B. leachi* from New Zealand (76). Alphaproteobacteria were found as dominant microbial members of *C. intestinalis* from Germany (77). Alphaproteobacteria and Gammaproteobacteria have also been shown to be highly abundant in previous studies of other tunicates (78), marine sponges (11, 79), corals (13, 80) and marine tube worms (15, 81). Bacteria from the *Rhodobacterales, Vibrionales, Oceanospirillales*, of class Alpha- and Gamma-proteobacteria, respectively, were previously isolated from *B. schlosseri* from the Yellow Sea, China (82). Although our results are consistent with previous findings, it remains unknown whether members of *B. schlosseri* microbial communities are permanent or transient, or if they are associated with specific tunicate body parts.

### 3.3 B. schlosseri metabolomes are distinct from surrounding seawater

To generate metabolomes of *B. schlosseri* samples and those of surrounding seawater, we used MPACT (45) to filter and prioritize reproducible features from our MS data, resulting in a total of 404 molecular features across all samples, with 16% of features unique to *B. schlosseri*, 35% unique to seawater, and 49% shared between both (Fig. S3). This high percentage of shared features is consistent with *B. schlosseri* as a filter-feeder of the surrounding seawater. Regardless of their shared features, hierarchical clustering (Fig. 3A) revealed two distinct clades for *B. schlosseri* and seawater, suggesting distinct metabolomic profiles. Interestingly, the *B. schlosseri* clade was further divided into three subclades, with subclade 2 including only one sample, distinct from subclades 3 and 4. This metabolomic variance amongst *B. schlosseri* samples, all collected at the same time and location, might suggest that microenvironment variation plays a role in the chemical ecology of *B. schlosseri*. The distinct metabolomic profiles of *B. schlosseri* and surrounding seawater were further confirmed using NMDS discriminant analysis (Fig. 3B), in which the overall metabolomic profiles were determined to form distinct groups.

**Fig. 3.**
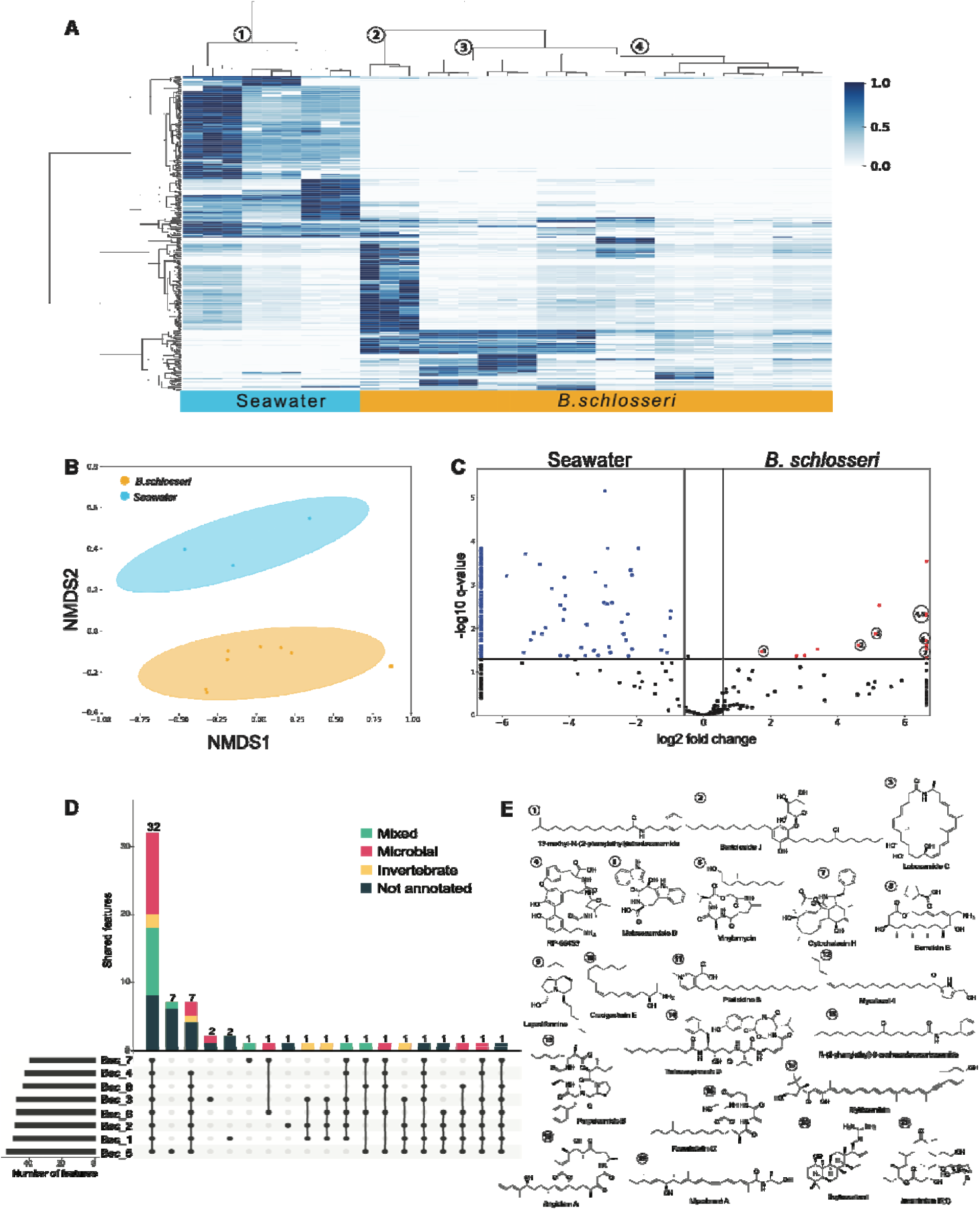
*B. schlosseri* and surrounding seawater have distinct metabolomes. (A) Heatmap of metabolomic features with the x-axis representing samples clustered by overall similarity using hierarchical clustering, and the y-axis representing features clustered by relative abundance across samples. Seawater and *B. schlosseri* were determined to separate into distinct subclades, with *B. schlosseri* further separating into three additional subclades. (B) Nonmetric multidimensional scaling (NMDS) analysis resulted in distinct clustering for *B. schlosseri* and seawater metabolomes (circles represent 95% confidence intervals; PERMANOVA q-value = 0.05, pseudo-F = 3.89). (C) Volcano plot comparing differential features from *B. schlosseri* and seawater, resulted in prioritization of several features that were significantly more abundant in *B. schlosseri* samples. Comparison of MS1 and MS2 data resulted in putative identification of several features (E). (D) Upset plot representing the *B. schlosseri* pan-metabolome, in which numbers above each bar indicate the total features and colors represent the proportion of annotated features, including their putative origin as reported in literature or from databases used for annotation. Metabolites of putative microbial origin are in pink, with those of putative invertebrate origin in yellow. Metabolites of mixed origin with matches to microbial and invertebrate metabolites are in green and those without annotation are dark blue. Interestingly, there are numerous features in the core metabolome (metabolites found in all samples) that are likely of microbial origin, perhaps representing compounds from consistently present microbes that may represent commensal species.

To explore which features varied most between *B. schlosseri* and seawater, we constructed a volcano plot (Fig. 3C) in which we observed approximately 15 features with significantly higher abundance in *B. schlosseri* than in seawater, and over 60 features with higher abundance in seawater. Thus, despite sharing almost half of their metabolomic features, significant variations in abundance of specific features may be indicative of the potential ecological roles of these metabolites in *B. schlosseri* or seawater. For example, highly abundant metabolites in *B. schlosseri* may be indicative of compound sequestration or accumulation (83, 24), whereas the high diversity of metabolites in seawater may be reflect the numerous chemical interactions and metabolic processes in seawater (9).

#### 3.3.1 B. schlosseri metabolome composed of metabolites from diverse sources

A total of 189 features of the *B. schlosseri* and seawater metabolomes were annotated using molecular weight, formula, and/or MS fragmentation. Of the total, 56 annotated features were found only in seawater, and 39 were found only in *B. schlosseri* (Fig. S3). Of the annotated features, 78% were reported from microbial sources, including bacteria, cyanobacteria, and fungi, while 7% were reported from marine invertebrates, including octocorals, sponges, and tunicates. We categorized 15% of the annotated features as deriving from mixed source, i.e. any feature that showed matches to both microbial and invertebrate metabolites.

We then sought to further explore annotations matching metabolites isolated from microbes. From our seawater samples, 95% of the annotated features were reported from microbial sources, such as porpoisamide (**13**) from the cyanobacteria *Lyngbya* sp. (84), thalassospiramides (**14**) from *Thalassospira* sp. (Alphaprotoebacteria; (85), and juvenimicin (**21**) from *Micromonospora chaela* (Actinobacteria; (86). In *B. schlosseri*, 46% of annotated features were from microbial sources, including from Actinobacteria, Myxobacteria, and Cyanobacteria. Three notable examples include RP-66453 (**4**) from *Streptomyces* spp. (87), lobosamide (**3**) from *Micromonospora* spp. (88), and myxalamid (**19**) and angiolam (**18**) from *Myxoccocus xanthus* and *Angioccocus disciformis*, respectively (89, 90). Interestingly, many of the annotated microbial metabolites from *B. schlosseri* were previously reported from microbial taxa that were found only in minor proportion in *B. schlosseri* (e.g., Deltaproteobacteria from Fig. 2, Cyanobacteria, and Actinobacteria), and we did not see annotated features previously reported from *B. schlosseri*’s most abundant microbial taxa, perhaps because these metabolites have yet to be identified or perhaps reflecting bias towards metabolically rich bacteria (e.g., Actinobacteria and Cyanobacteria) in natural product databases.

We then considered annotations from our samples matching metabolites previously isolated from invertebrates. Unfortunately, no definitive matches were found to botryllin, the only previously reported metabolite from the *Botryllus* genus. However, several of our annotated metabolites matched those reported from other invertebrate sources, including sponges, soft corals, and other tunicates. For example, putative annotations included the metabolites lepadiformine (**9**) and crucigasterin (**10**), which were initially isolated from the colonial tunicates *Clavelina lepadiformis* and *Pseudodistoma crucigaster,* respectively (91, 92). Another annotated invertebrate metabolite included mytiloxanthin (**17**), originally isolated from the sea mussel *Mytilus californianus* (93), but also reported from the ascidian *Halocynthia roretzi* (94). Other matches to metabolites from microbe-associated invertebrates included *N*-(phenylethyl)-9-oxohexadecacarboxamide (**15**) from the octocoral *Telesto riisei* (95) and mycalazol 4 (**12**) from the sponge *Mycale micracanthoxea* (96).

Tunicates have been recognized for their prolific production of biologically relevant metabolites (25–27, 97), although many studies have revealed that many of these interesting metabolites may be produced by associated bacteria (83, 24, 98). Our findings of both microbial- and invertebrate-derived metabolites from within our *B. schlosseri* samples are consistent with previous reports (99, 100, 82, 101). In addition, several of the microbial metabolites annotated from our *B. schlosseri* samples have been reported to have cytotoxic activity (102, 103), suggesting *B. schlosseri* may have adapted to harbor associations with microorganisms capable of producing defensive compounds. Additional data related to the microbes and their metabolites that are consistently associated with *B. schlosseri* will allow for more definitive understanding of *B. schlosseri* microbiomes.

#### 3.3.2 B. schlosseri *pan-metabolome*

Genomics research has been using a pan-genome approach to compare the shared and unique genomic components of related bacterial species (104, 105). More recently, the pan-genome concept has been applied to complex microbiome samples (106–108), to differentiate shared or unique microbial strains amongst microbiome members. We sought to apply similar concepts to our *B. schlosseri* metabolomics data, given the complexities of host-associated microbial communities, especially marine filter feeders, such as sponges and tunicates. By classifying metabolomic features in the context of a pan-metabolome, we considered the whole *B. schlosseri* metabolome as a unique entity with the host and associated microbiota (i.e. holobiont) that may form symbiotic associations, contributing to each other’s defense, nutrition, protection, immunity, and/or development. Thus, we introduce the concept of a pan-metabolome to classify metabolites according to their presence or absence across complex microbiome samples. Under this premise, the core metabolome includes metabolites found in all samples, while the flexible metabolome includes those found only in some samples. Metabolites from the core metabolome would be presumed to be highly important for the biology of the system, regardless of source, whereas metabolites from the flexible metabolome might be associated to an organism’s environment or might be only transiently associated.

Among the 64 metabolomic features found in our *B. schlosseri* samples, 32 were shared across all eight samples (Fig. 3D), comprising their core metabolome. In contrast, five of the eight *B. schlosseri* samples contained features found only in that sample, one of which had seven unique features while the other four each had either one or two unique features. The remaining three *B. schlosseri* samples shared their features with at least one other sampled colony (Fig. 3D). Those features not found in all eight samples comprise the flexible metabolome.

We were able to annotate several features in both the core and flexible metabolomes. Surprisingly, we found more features related to microbial sources in the core metabolome than in the flexible metabolome. For example, the actinomycete-derived metabolites lobosamide C (**3**) and RP-66453 (**4**) were annotated in the core metabolome, which is interesting because although members of the Actinobacteria have been reported to be tunicate-associated, they usually occur in lower abundance than members of Proteobacteria (109). We also observed the fungal metabolite cytochalasin (**7**), produced by multiple members of the phylum Ascomycota, which has previously been reported as an important tunicate-associated phylum (110, 24).

Several features from mixed sources were annotated in the core metabolome, including from sponges, corals, and fungal metabolites. Crucigasterin (**10**) from the tunicate *Pseudodistoma crucigaster* (92) was found in the core metabolome, as well as fusaristatin (**16**), which has been reported from the octocoral *Eunicea fusca* and its associated fungus *Pithomyces* sp. (111). Within the flexible metabolome, seven annotated features were from microbial sources, including angiolam (**18**), scytoscalarol (**20**), and borrelidin (**8**), all discussed above. Four features were from invertebrate sources, including lepadiformine (**9**) (91) found in seven of our *B. schlosseri* samples, as well as platisidine (**11**) (112), mycalazol (**12**), and *N*-(phenylethyl)-9-oxohexadecacarboxamide (**15**) (95) all three of which were found in half of our *B. schlosseri* samples.

Additional sampling, across numerous locations and various environments, will be needed to confidently define members of the *B. schlosseri* pan-metabolome. However, even with our current sample size, several interesting patterns in metabolite distribution were observed leading to the question of whether the microbial taxa producing *B. schlosseri* core metabolites might be in symbiotic association.

### 3.4 Integrating the metallome, microbiome, and metabolome of B. schlosseri and surrounding seawater

To further investigate how metals, microbes, and metabolites interact within *B. schlosseri* and surrounding seawater, we used an *N*-integration model (54) to determine multi-omics correlations. The resulting individual (s)PLS-DA analyses for metallome, microbiome, and metabolome (Fig. 4A) indicated that *B. schlosseri* and seawater represent separate groups. Interestingly, we observed tight clustering of seawater metals and moderately tight clustering of *B. schlosseri* metabolites, indicating these groups were relatively similar for those particular - omics analyses.

**Fig. 4.**
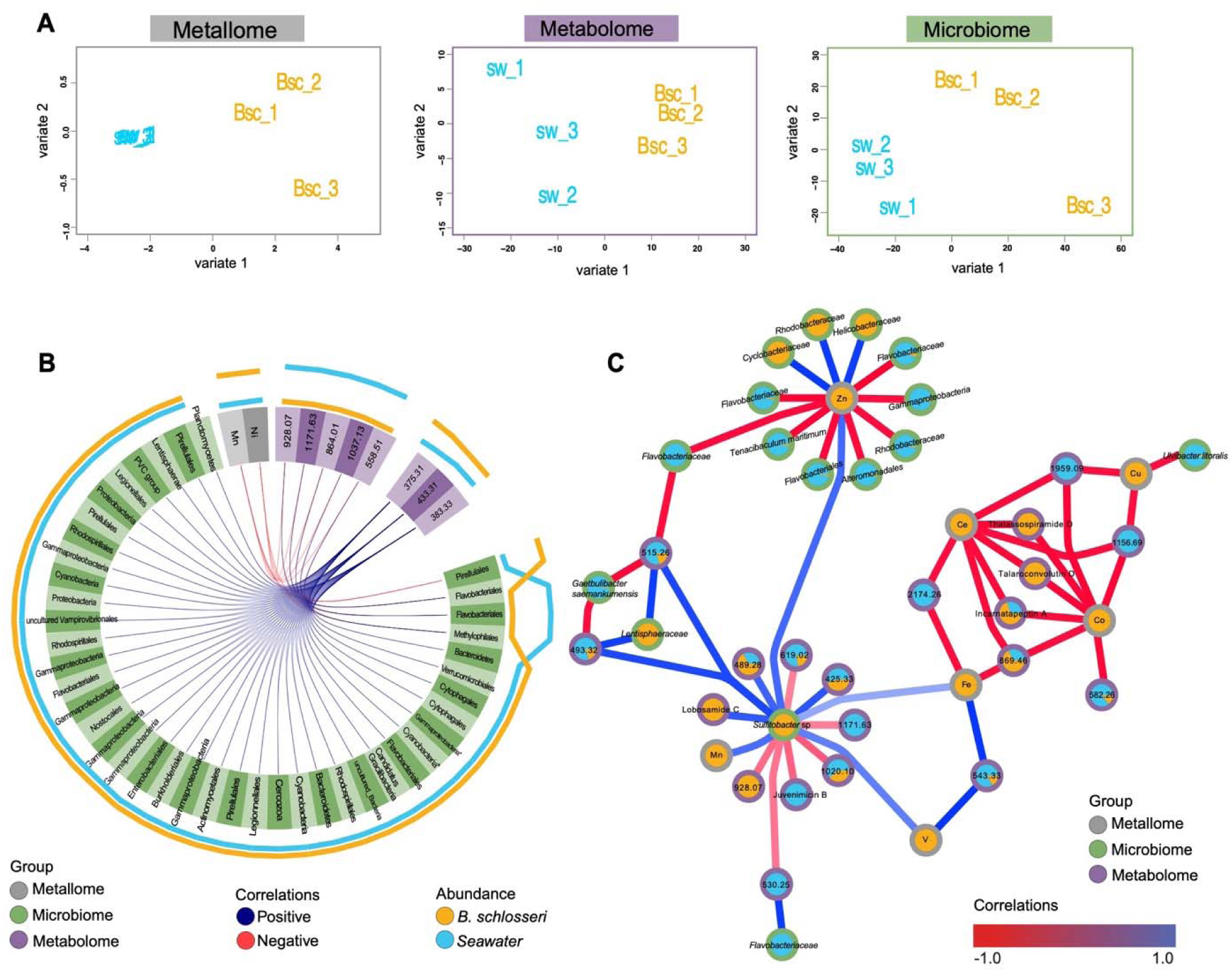
Multi-omic integration model revealed significant interactions between metals, microbes, and metabolites in *B. schlosseri* and surrounding seawater. (A) Individual (s)PLS-DA analyses for metallome, microbiome, and metabolome from the tuned Diablo model (M-fold validation, with 3 folds and 10 iterations) indicated that *B. schlosseri* (yellow, n = 3) and seawater (blue, n = 3) represent separate groups for each of the individual -omics analyses. (B) Using a circos plot built from Spearman pairwise correlations, numerous significant correlations (r > 0.9) were observed amongst members of the metallome (grey), microbiome (green), and metabolome (purple), with blue connections representing positive correlations and red connections representing negative correlations. The outer lines represent the abundance of each feature in *B. schlosseri* (yellow) and seawater (blue). (C) Network visualization of correlations from the DIABLO *N*-integration model of select multi-omics features (full network in Fig. S4). Using this integrated model, numerous intriguing relationships were apparent between specific metals, microbes, and metabolites. Node borders indicate the -omics analysis (grey for metallome, green for microbiome, purple for metabolome). Node pie charts indicate the proportion of features from *B. schlosseri* (yellow) or seawater (blue). Nodes are labelled with either the lowest microbial taxonomic resolution, metal element symbol, putative metabolite annotation, or mass in Daltons.

Next, we sought to further integrate our multi-omics data using a circos plot to visualize both positive and negative correlations between specific features that might be driving the model (Fig. 4B). For this analysis, features with directly proportional abundances (e.g., high abundance of a specific metal correlated to high abundance of a specific metabolite; or low abundance of a specific microbe correlated to low abundance of a specific metal) were denoted as positive correlations, while inverse trends in abundances between features (high versus low or low versus high) were denoted as negative correlations. Interestingly, although iron, manganese, and zinc were the most abundant metals in our *B. schlosseri* and seawater samples (Fig. 1A), in the reduced model (tuned mode) used to generate the circos plot, manganese and nickel were the metals with the most significant interactions with members of the metabolome and microbiome (Fig. 4B). The most significant correlations for manganese and nickel were all negative correlations with five unannotated metabolites, ranging from 558.5 Da to 1171.6 Da, all of which were more abundant in seawater than in *B. schlosseri*. In addition, both manganese and nickel had significant negative correlation with an ASV from the order *Pirellulales*, thus as manganese and nickel increased, *Pirellulales* decreased, which might indicate sensitivity of this strain to manganese and nickel. The circus plot also revealed strong positive correlations between three unannotated features (masses of 375.3 Da, 383.3 Da and 433.3 Da), all enriched in the *B. schlosseri* metabolome, with many microbial taxa, including members from the phyla *Proteobacteria*, *Actinobacteria*, and *Bacteroidota*.

To extend beyond pairwise correlations, we used networking analysis to visualize the *N*-integration model multi-omics correlations (Fig. 4C), revealing numerous interactions between metals, microbes, and metabolite. The metals, cerium and cobalt, were highly negatively correlated to several metabolites, including thalassopiramide D, talaroconvolutin D and incarnatapeptin A, within *B. schlosseri* tissue. Although none of these metabolites were directly associated with iron in our study, the thalassospiramide producing *Thalassospira* spp. are known iron reducing bacteria (113), and talaroconvolutin D is a fungal compound that induces ferroptosis by ROS upregulation (114). Copper was also found to be negatively correlated with several metabolites, including those with masses 1959.1 and 1156.6 Da, and was also negatively correlated to *Ulvitobacter litoralis*. Copper is an important nutrient in the marine environments at trace concentrations, especially for primary producers (115), however it has been observed that long exposure of *B. schlosseri* colonies to high concentrations of copper (e.g., 5 mg and 48 hrs) can be lethal (75). Additional experiments are needed to explore the role of this micronutrient in our system.

In our model (Fig. 4C), zinc exhibited negative and positive correlations with many microbial taxa, including members of Gammaproteobacteria, Flavobacteriales, and Alphaproteobacteria (Rhodobacteriales). Notably, the fish pathogen *Tenacibaculum maritimum* (Flavobacteriales), present in much higher proportion in seawater than *B. schlosseri* tissue, was strongly negatively correlated with zinc. In contrast, a member of the family *Helicobacteraceae* (Epsilonproteobacteria) was positively correlated with zinc and was more abundant in *B. schlosseri* than in sweater. Previous research has shown that marine members of *Helicobacteraceae* have been found in commensal relationships with gastropods, as well as in association with the coelomic fluid in echinoderms (116, 117), suggesting they might be beneficial host-associated bacteria, although their ecological role has not been fully described.

Iron and vanadium, both more abundant in *B. schlosseri*, were both positively correlated with a member of the genus *Sulfitobacter*, which was also more abundant in *B. schlosseri* and which had further correlations with several metabolites and with manganese (Fig. 4C). Iron and vanadium also exhibited positive correlations with an unannotated metabolite (mass 543.3 Da) that was more abundant in seawater. Previous research has shown bioaccumulation of iron in hemocytes from *B. schlosseri* embryos, suggesting an important role of iron the immune system and development of *B. schlosseri* (118). Interestingly, tunicates are known to sequester vanadium to a very high concentrations (119, 120), and it has been suggested that vanadium can act as a feeding deterrent for fish (121). In addition, although not well understood, there are suggestions that symbiotic bacteria might be involved in vanadium accumulation (122). For example, previous studies have found the bacterial genera, *Pseudomonas* and *Ralstonia*, in high abundance in vanadium-rich ascidians, particularly in the pharynx, which is involved in vanadium absorption (123). *Sulfitobacter* species are sulfite-oxidizing bacteria that have been previously shown to act as pathogens, inducing cell death of marine microalga (124), although their role in our system remains unclear especially since both the *Sulfitobacter* sp. and essential metals such as iron and manganese were all more abundant in *B. schlosseri*.

Many of the metals in our multi-omics integration model are essential nutrients for hosts and bacteria, including manganese, iron, cobalt, nickel, copper, and zinc, some of which have been reported to be important in forming pathogenic or symbiotic interactions with eukaryotic host cells (125). For example, in mammalian cells, the immune system inhibits bacterial growth by withholding critical metals, including manganese, iron, and zinc to starve invading bacteria, resulting in nutritional immunity (126, 127). In other eukaryotic hosts, there is also evidence of increased concentrations of copper, zinc, iron, and manganese in host defenses against bacteria, especially during colonization (128–130). Additional experiments are needed to further explore the ecological roles that associated bacterial taxa and metabolites may have in the context of *B. schlosseri* metal binding, accumulation, and transport.

## 4. Conclusion

Herein, we compared the microbiome, metals, and metabolites found in *B. schlosseri* with those in the surrounding seawater. We identified taxa within the microbial community of *B. schlosseri* that were either more abundant than in seawater (e.g., *Gammaproteobacteria*, *Episilonproteobacteria*) or less abundant (e.g., *Alphaproteobacteria*, *Flavobacteriia*), providing indications that *B. schlosseri* employs some mechanism(s) for microbiome selection. Similarly, our baseline metallome studies demonstrated that *B. schlosseri* sequesters several metals with a substantial fold-increases over those found in seawater. We also found several microbial metabolites to have increased abundance in *B. schlosseri* with those in seawater, including some that may bind metals based their chemical structures. We introduced the concept of a pan-metabolome to classify metabolites as of high importance based on their presence or absence across samples, finding that *B. schlosseri* samples had a high proportion of shared metabolites, indicating a closed pan-metabolome. Finally, we integrated our multi-omics data to investigate which microbial taxa and metabolites co-vary with metals, finding several that are positively or negatively correlated. These results suggest that metals and metabolites play important roles in structuring the *B. schlosseri* microbiome, making it an ideal test system for future mechanistic investigation and for unraveling possible symbiotic relationships based on metabolite interactions with microbes and metals.

This study is the first multi-omics approach connecting the metallome to the microbiome and metabolome of *B. schlosseri*. Tunicates, including *B. schlosseri*, are integral members of the marine environment, providing important, bioactive natural products, with recent interest in the role of their associated microbes. However, while some tunicate taxa have been studied for their ability to bioaccumulate trace metals (29, 30), the microbial role in this process, and that of associated metabolites, remains poorly understood. Our study bridges the knowledge gap surrounding the intricate relationships among organisms, their symbionts, the metabolites they produce, and the sequestered and environmental metals. The correlations we observed reveal the important interactions between metals, microbes, and metabolites in *B. schlosseri*, providing a baseline for future studies of metal accumulation from a host-microbiome context. Moreover, this research establishes a foundation for comprehending the ecological importance of trace metals in host-microbe systems, potentially paving the way for the discovery of new natural products with unique therapeutic applications.

## Acknowledgements

Funding for this work was provided by the University of Connecticut (UConn) Convergence Awards for Research in Interdisciplinary Centers (CARIC; to MJB, SVN, AAB) and from the National Science Foundation Graduate Research Fellowships Program (GRFP; to CMD and MH). We also thank Dr. Kendra Maas from the UConn Microbial Analysis, Resources and Services (MARS) facility for assistance with amplicon library preparation and sequencing. Thanks are due to the UConn Center for Environmental Sciences and Engineering, especially Christopher Perkins and Sneiguole Stapcinskaite, for assistance with inductively coupled plasma-mass spectrometry (ICP-MS).

## Declaration of competing interest

The authors do not have any competing interest to declare.

## CRediT authorship contribution statement

**D. Guillén Matus, C. Donaghy, N. Vijayan A. Angeles-Boza, S. Nyholm, M. Balunas:** designed and performed research, analyzed data, and wrote the paper. **Z. Lane:** performed research, contributed to writing paper. **M. Howell, G. Glavin:** performed research. All authors edited the final manuscript.

## Data availability

Sequencing data were deposited to the National Center for Biotechnology Information (NCBI) GenBank (https://www.ncbi.nlm.nih.gov/genbank/) under accession numbers PRJNA1039848. Metabolomics data were deposited to the Mass Spectrometry Interactive Virtual Environment repository MassIVE (https://massive.ucsd.edu/ProteoSAFe/static/massive.jsp) under accession numbers MSV000096313 (ftp://massive.ucsd.edu/v07/MSV000096313/). Code is available on GitHub (https://github.com/BalunasLab/Botryllus-schlosseri-multiomics).

## Appendix A. Supplementary data

Supplementary data for this article can be found online at https://doi.org/10.1028/msystems.xxx.

## Notes

### Competing Interest Statement

The authors have declared no competing interest.

### Summary of Updates

Provide additional details and revise figures

