## Supplementary Materials for "Multi-omics Analysis Reveals Important Role for Microbial-derived Metabolites from *Botryllus schlosseri* in Metal Interactions"

**This document contains:**

Table S1. Limits of detection (LODs) for metals from tunicate and seawater samples.

Table S2. Raw and mean metal concentrations [± standard deviation (SD)] for tunicate and seawater samples acquired using ICP-MS with p-values determined using two-tailed, homoscedastic t-tests.

Table S3. Positive correlations of metals, metabolites and bacterial ASVs in *B. schlosseri*, based on Diablo multi-omics model.

Table S4. Negative correlations of metals, metabolites and bacterial ASVs in *B. schlosseri*, based on Diablo multi-omics model.

Table S5. Assessment of putative annotations for important metabolites in *B. schlosseri* system, including those found to correlate with metals, bacterial taxa, or found in the pan-metabolome.

Figure S1. Rarefaction curves of observed amplicon sequence variants (ASVs) from 16S rRNA gene sequences for microbiota samples at a sequencing depth of 4,173 reads.

Figure S2. Relative abundance of microbiota. Bar graph of relative abundance of (A) bacterial communities in *B. schlosseri* and seawater and (B) bacterial taxa of core ASVs of *B. schlosseri.*

Figure S3. Overview of the distribution of features across *B. schlosseri* and surrounding seawater samples, including the proportion of identified features and their biological sources based on databases used for metabolite annotation.

Figure S4. Multi-omics correlation network showing all interactions from the Diablo full integration model.

Figure S5. First component of pairwise Pearson correlations from Diablo multi-omics integration model for *B. schlosseri* and the surrounding seawater.

Figure S5. Second component of pairwise Pearson correlations from Diablo multi-omics integration model for *B. schlosseri* and the surrounding seawater.

Table S1. Limits of detection (LODs) for metals from tunicate and seawater samples.

| | **Ce** | **V** | **Fe** | **Mn** | **Co** | **Ni** | **Cu** | **Zn** |
|---|---|---|---|---|---|---|---|---|
| **Tunicate LOD (ppm)** | 0.1 | 0.1 | 13 | 0.1 | 0.1 | 0.1 | 0.1 | 3 |
| **Seawater LOD (ppm)** | 0.002 | 0.001 | 0.001 | 0.001 | 0.001 | 0.001 | 0.001 | 0.001 |

Table S2. Raw and mean metal concentrations [± standard deviation (SD)] for tunicate and seawater samples acquired using ICP-MS with p-values determined using two-tailed, homoscedastic t-tests.

| | **Ce (ppm)** | **V (ppm)** | **Fe (ppm)** | **Mn (ppm)** | **Co (ppm)** | **Ni (ppm)** | **Cu (ppm)** | **Zn (ppm)** |
|---|---|---|---|---|---|---|---|---|
| ***B. schlosseri*** | 3.10 | 5.29 | 1928.89 | 79.44 | 0.76 | 6.16 | 11.98 | 36.88 |
| ***B. schlosseri*** | 4.30 | 8.01 | 2776.28 | 98.66 | 0.87 | 8.02 | 12.17 | 74.88 |
| ***B. schlosseri*** | 5.07 | 7.56 | 2877.61 | 89.05 | 1.12 | 13.32 | 17.16 | 58.31 |
| ***B. schlosseri* - mixture** | 3.19 | 5.29 | 2094.25 | 83.69 | 0.82 | 6.89 | 12.57 | 40.99 |
| **Tunicate Mean** | 3.9 ± 0.9 | 6.5 ± 1.5 | 2419.3 ± 477.4 | 87.7 ± 8.3 | 0.9 ± 0.2 | 8.6 ± 3.2 | 13.5 ± 2.5 | 52.8 ± 17.4 |
| **Seawater** | 0.00136 | < LOD | 0.02851 | < LOD | < LOD | 0.00394 | < LOD | 0.00356 |
| **Seawater** | 0.00235 | < LOD | 0.55626 | 0.00923 | 0.00172 | 0.07132 | 0.00298 | 0.00548 |
| **Seawater** | 0.00215 | < LOD | 1.071 | 0.02083 | 0.00303 | 0.13087 | 0.00584 | 0.00325 |
| **Seawater Mean** | 0.0020 ± 0.0005 | below LOD | 0.6 ± 0.5 | 0.015 ± 0.008 | 0.0024 ± 0.0009 | 0.07 ± 0.06 | 0.004 ± 0.002 | 0.004 ± 0.001 |
| **p-value** | 0.002 | - | 0.0004 | 0.0001 | 0.002 | 0.007 | 0.002 | 0.004 |

Table S3. Positive correlations of metals, metabolites and bacterial ASVs in *B. schlosseri*, based on Diablo multi-omics model.

| **Metal** | **Metabolite** | **Bacteria** |
|---|---|---|
| Iron | 543.3 Da | |
| | | *Sulfitobacter* |
| Vanadium | 543.3 Da | |
| | | *Sulfitobacter* |
| | 375.3 Da | *Proteobacteria* |
| | | *Actinobacteria* |
| | | *Bacteroidetes* |
| | 383.3 Da | *Proteobacteria* |
| | | *Actinobacteria* |
| | | *Bacteroidetes* |
| | 433.3 Da | *Proteobacteria* |
| | | *Actinobacteria* |
| | | *Bacteroidetes* |

Table S4. Negative correlations of metals, metabolites and bacterial ASVs in *B. schlosseri*, based on Diablo multi-omics model.

| **Metal** | **Metabolite** | **Bacteria** |
|---|---|---|
| manganese | | *Pirellulales* |
| | 558.51 Da | |
| | 864.01 Da | |
| | 928.07 Da | |
| | 1037.13 Da | |
| | 1171.63 Da | |
| nickel | 558.51 Da | |
| | 864.01 Da | |
| | 928.07 Da | |
| | 1037.13 Da | |
| | 1171.63 Da | |
| cerium | thalassopiramide D | |
| | talaroconvolutin D | |
| | incarnatapeptin | |
| cobalt | thalassopiramide D | |
| | talaroconvolutin D | |
| | incarnatapeptin | |
| copper | 1959.1 Da | |
| | 1156.6 Da | |
| zinc | | *Tenacibaculum maritimum* (Flavobacteriales) |

Table S5. Assessment of putative annotations for important metabolites in *B. schlosseri* system, including those found to correlate with metals, bacterial taxa, or found in the pan-metabolome.

| *m/z* | RT (min) | Putative annotation | Molecular Formula | MS identification level of confidence* |
|---|---|---|---|---|
| 346.31061 | 5.90 | 13-methyl-*N*-(2-phenylethyl)tetradecanamide | $C_{23}H_{39}NO$ | Level 3 |
| 599.40824 | 7.21 | bartoloside J | $C_{34}H_{59}ClO_6$ | Level 3 |
| 468.30795 | 6.02 | lobosamide C | $C_{29}H_{41}NO_4$ | Level 3 |
| 617.25738 | 5.72 | RP-66453 | $C_{33}H_{36}N_4O_8$ | Level 3 |
| 376.12976 | 5.02 | malassezindole B | $C_{21}H_{17}N_3O_4$ | Level 2 |
| 494.32403 | 6.15 | vinylamycin | $C_{26}H_{43}N_3O_6$ | Level 3 |
| 516.27187 | 6.03 | cytochalasin H | $C_{30}H_{39}NO_5$ | Level 3 |
| 494.34708 | 5.99 | borrelidin B | $C_{28}H_{47}NO_6$ | Level 3 |
| 294.27905 | 5.97 | lepadiformine | $C_{19}H_{35}NO$ | Level 2 |
| 280.26334 | 5.88 | crucigasterin E | $C_{18}H_{33}NO$ | Level 2 |
| 362.30536 | 5.97 | platisidine B | $C_{23}H_{39}NO_2$ | Level 3 |
| 376.32120 | 6.05 | mycalazol 4 | $C_{24}H_{41}NO_2$ | Level 2 |
| 621.36037 | 5.87 | porpoisamide B | $C_{33}H_{50}N_4O_6$ | Level 3 |
| 832.48931 | 5.88 | thalassospiramide D | $C_{46}H_{65}N_5O_9$ | Level 3 |
| 374.30580 | 6.11 | N-(2-phenylethyl)-9-oxohexadecacarboxamide | $C_{24}H_{39}NO_2$ | Level 3 |
| 482.32460 | 6.43 | fusaristatin C | $C_{25}H_{43}N_3O_6$ | Level 3 |
| 599.40824 | 7.21 | mytiloxanthin | $C_{40}H_{54}O_4$ | Level 2 |
| 610.37117 | 6.80 | angiolam A | $C_{34}H_{53}NO_7$ | Level 3 |
| 438.29767 | 6.94 | myxalamid A | $C_{26}H_{41}NO_3$ | Level 3 |
| 438.34184 | 7.17 | scytoscalarol | $C_{26}H_{45}N_3O$ | Level 3 |
| 568.38693 | 6.65 | juvenimicin B(1) | $C_{31}H_{53}NO_8$ | Level 3 |

*Schymanski's Rules for Using MS to Assess Confidence (Schymanski et al., 2014):
Level 1: confirmed structure by reference standard (MS, MS/MS, RT, Reference Standard)
Level 2: probable structure by a) library spectrum match b) diagnostic evidence (MS, MS/MS, library MS/MS, experimental data)
Level 3: Tentative candidate(s) structure, substituent, class (MS, MS/MS, experimental data)
Level 4: Unequivocal molecular formula (MS, isotope/adduct)
Level 5: Exact mass of interest (MS)

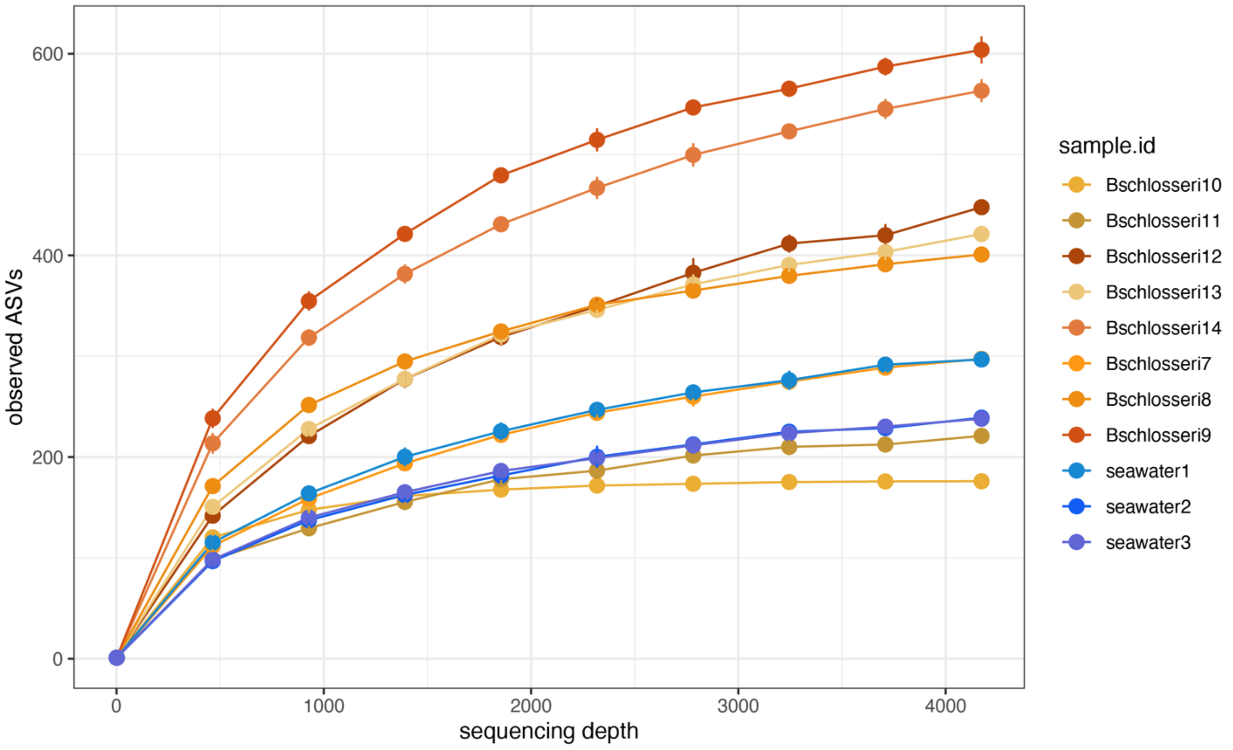


Figure S1. Rarefaction curves of observed amplicon sequence variants (ASVs) from 16S rRNA gene sequences for microbiota samples at a sequencing depth of 4,173 reads.

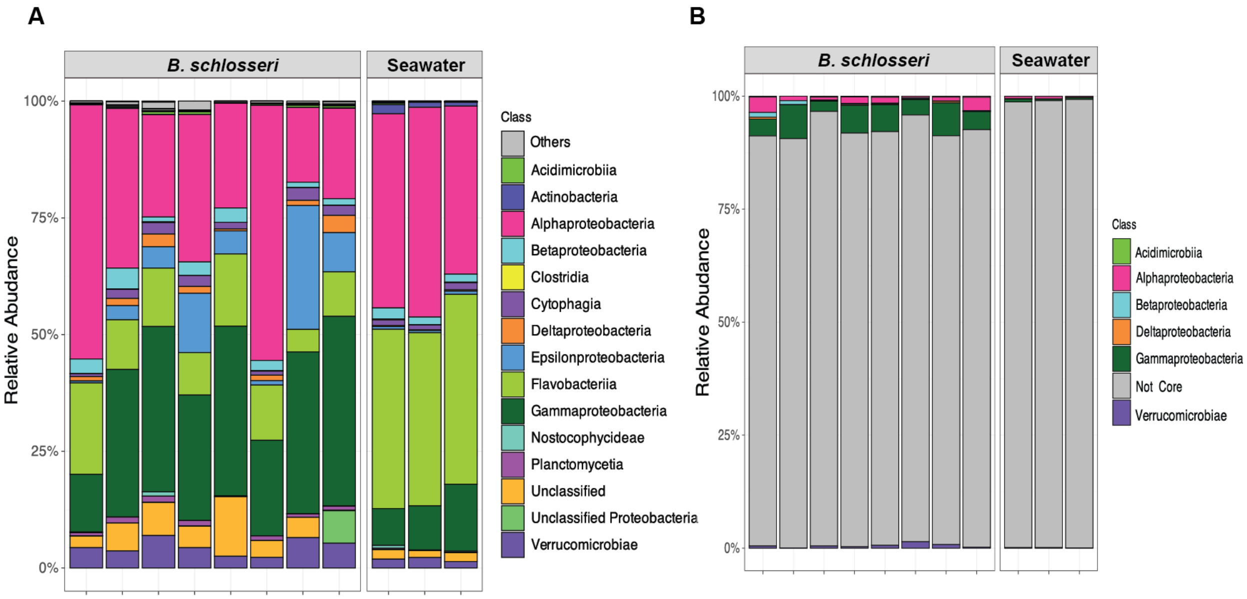


Figure S2. Relative abundance of microbiota. Bar graph of relative abundance of (A) bacterial communities in *B. schlosseri* and seawater and (B) bacterial taxa of core ASVs of *B. schlosseri.*

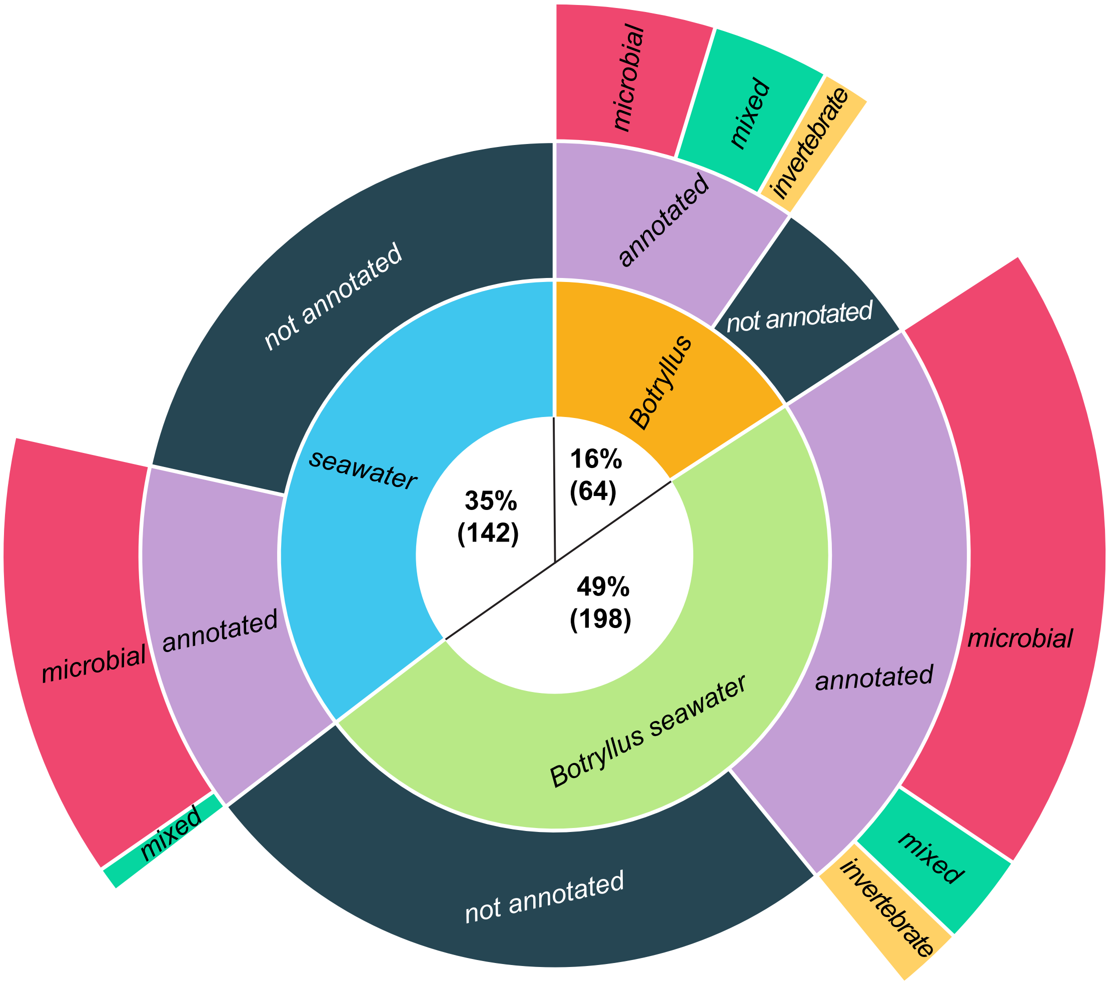


Figure S3. Overview of the distribution of features across *B. schlosseri* and surrounding seawater samples, including the proportion of identified features and their biological sources based on databases used for metabolite annotation.

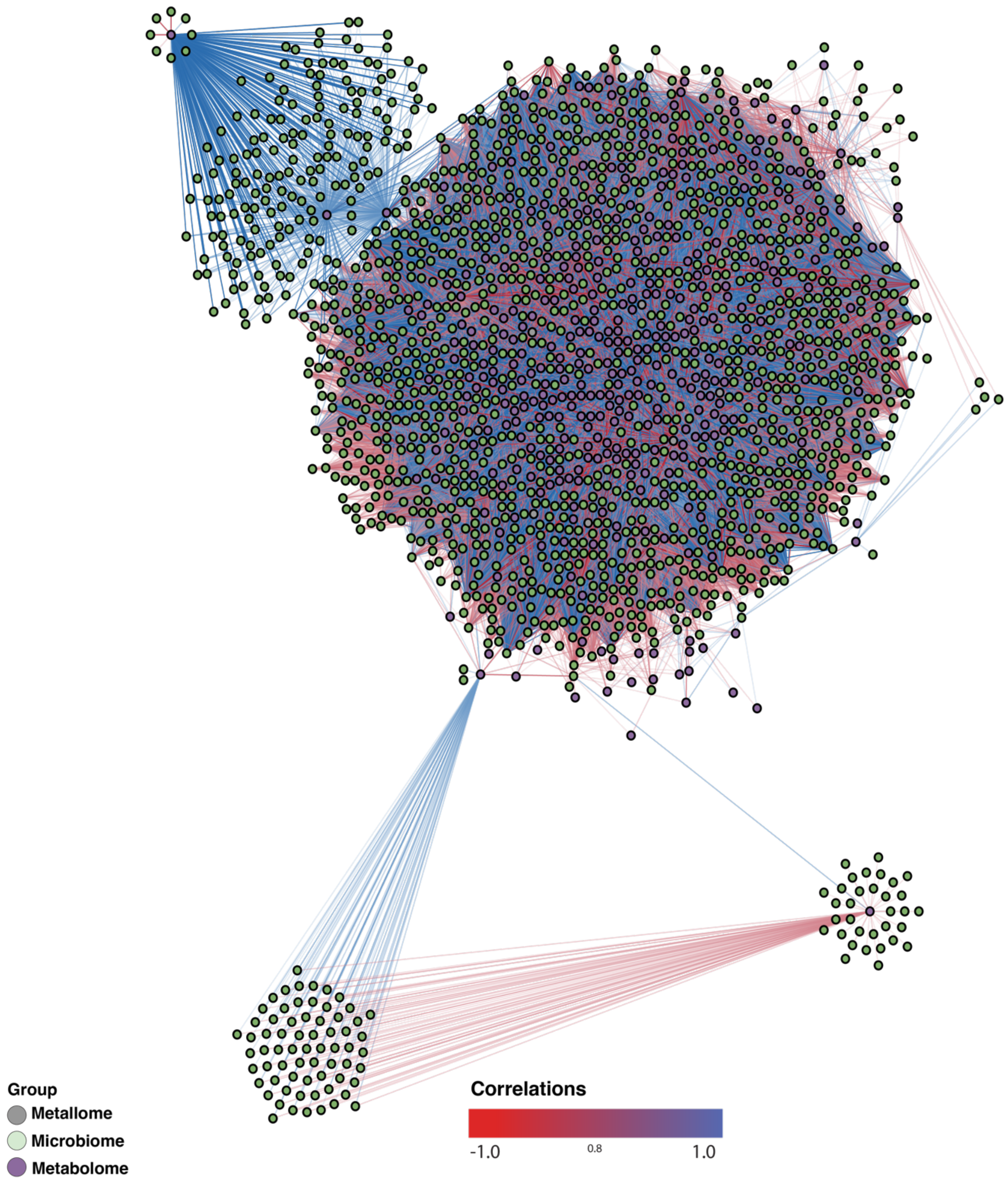


Figure S4. Multi-omics correlation network showing all interactions from the Diablo full integration model. Edge cutoff is 0.8 and -0.8 and the color represents negative (red) or positive (blue) correlations between members of the metallome, microbime, and metabolome.

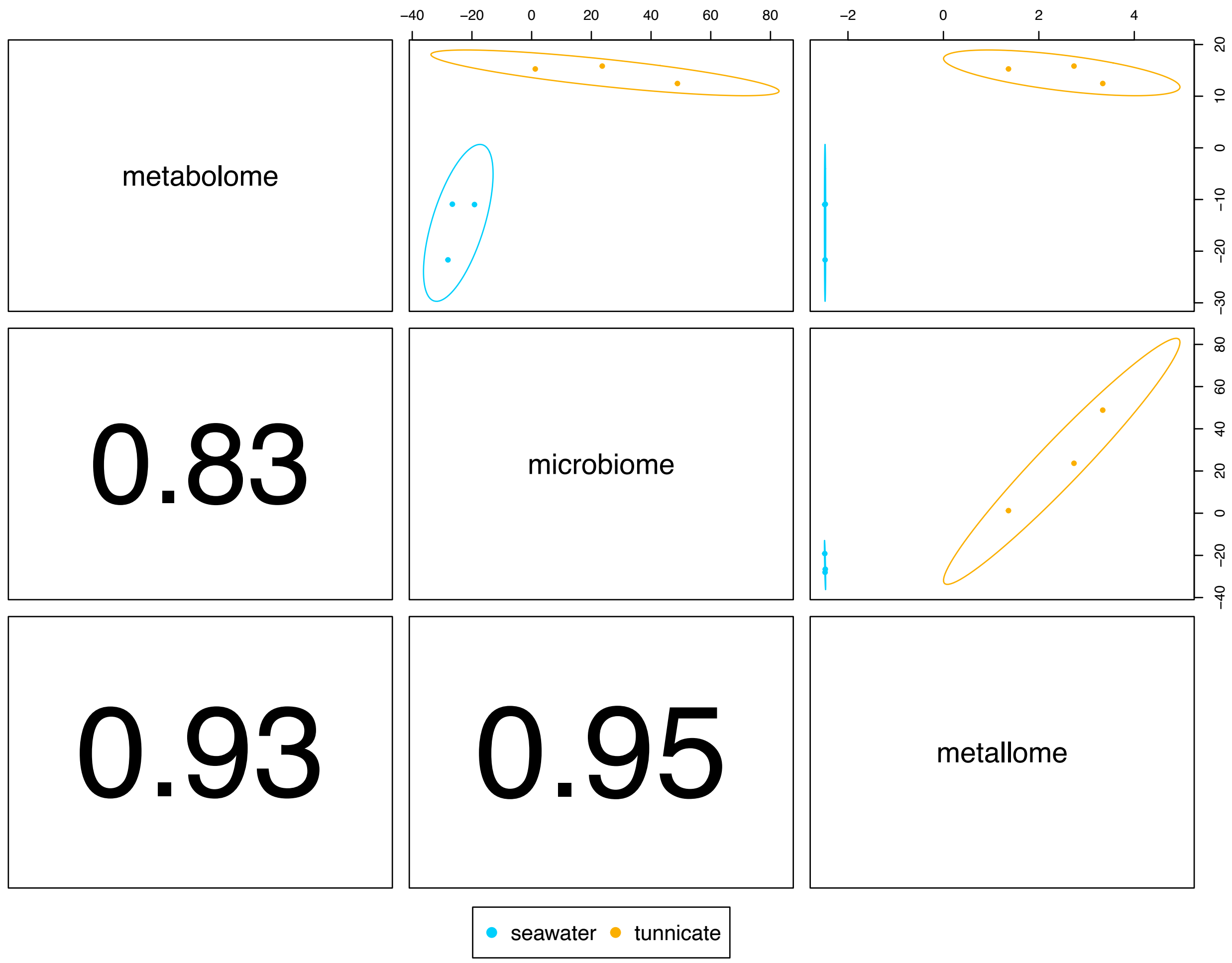


Figure S5. First component of pairwise Pearson correlations from Diablo multi-omics integration model for *B. schlosseri* and the surrounding seawater.

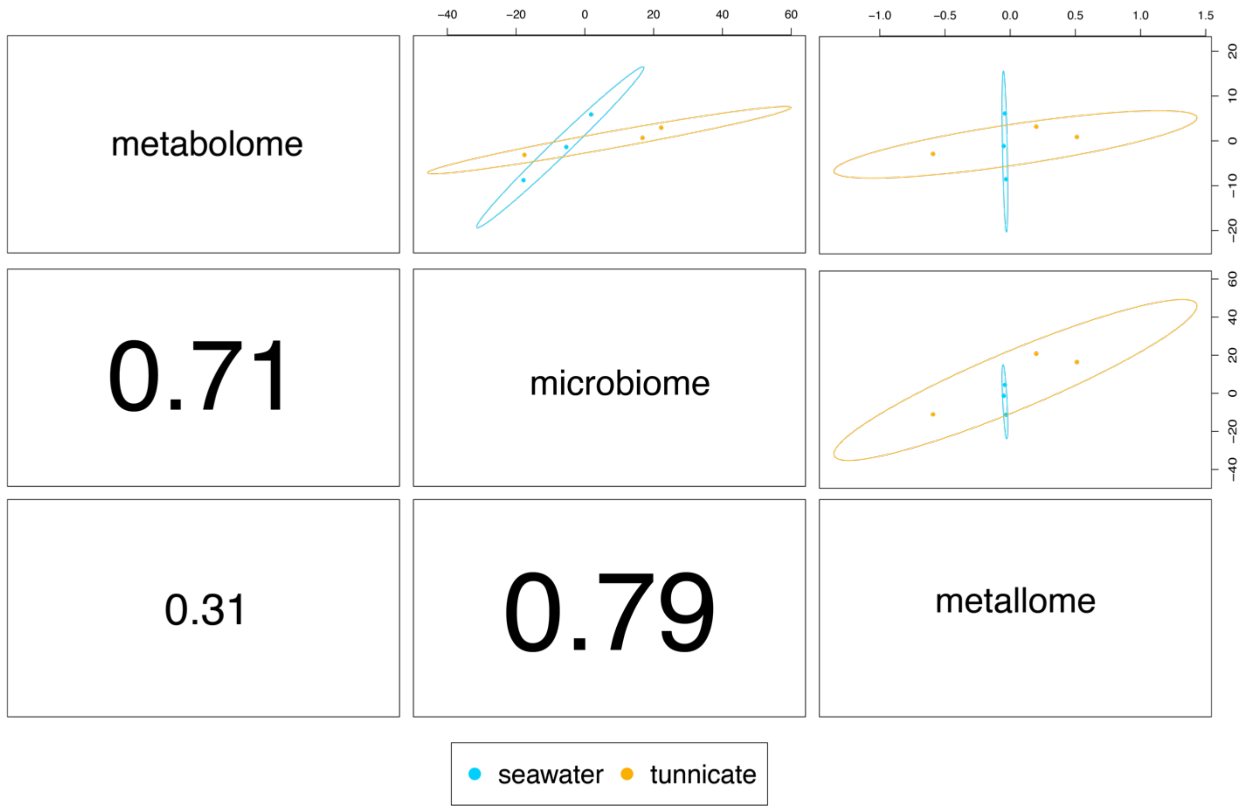


Figure S6. Second component of pairwise Pearson correlations from Diablo multi-omics integration model for *B. schlosseri* and the surrounding seawater.
